# A non-Lamarckian model for the inheritance of the epigenetically-coded phenotypic characteristics: a new paradigm for Genetics, Genomics and, above all, Ageing studies

**DOI:** 10.1101/492280

**Authors:** Xavi Marsellach

**Affiliations:** Independent Researcher, (L’ Hospitalet de Llobregat, Barcelona, Catalan Republic)

## 1. Introduction

In this paper I would like to introduce some theoretical considerations, which, if proven right by the experimental frameworks designed to test them, might stand for a paradigmatic change on genetics, genomics and ageing studies. This paper further develops a theoretical framework previously proposed by this author (Marsellach, 2017). Despite this, this paper is written as a standalone paper, which describe in more detail the previously hinted model (Marsellach, 2017), and adds emphasis in the putative implications that, the proposed theoretical framework, might have in describing the way in which the epigenetic information is transmitted from one generation to the following ones, and in the implications that this might have for genetics, genomics and ageing studies.

## 2. The Laws of Mendel, Schrödinger’s cat-like interpreted

The best way to describe the model for the inheritance of the epigenetically-coded phenotypic characteristics that I am proposing, is by asking you to: 1) do not trust Mendel (Mendel, 1866), and, at the very same time, to 2) do trust Mendel (Mendel, 1866). This leads inevitably to a Schrödinger’s cat-like scenario (see Figure 1). In other words, this implies that, while Mendelian inheritance is, almost, always true for the genetically-coded phenotypic characteristics (see Figure 1A), this is not the case for the epigenetically-coded phenotypic characteristics (see Figure 1B). As detailed later, in an ideal scenario, the Laws of Mendel should never be true for the epigenetically-coded phenotypic characteristics. This is due to the fact that there is a meiotic epigenetic repair program that should repair all the epimutations accumulated so far (see later and (Marsellach, 2017)). But the fact is that, sometimes, the Laws of Mendel, are, indeed, true for the epigenetically coded phenotypic characteristics (see below for details).

**Figure 1:**
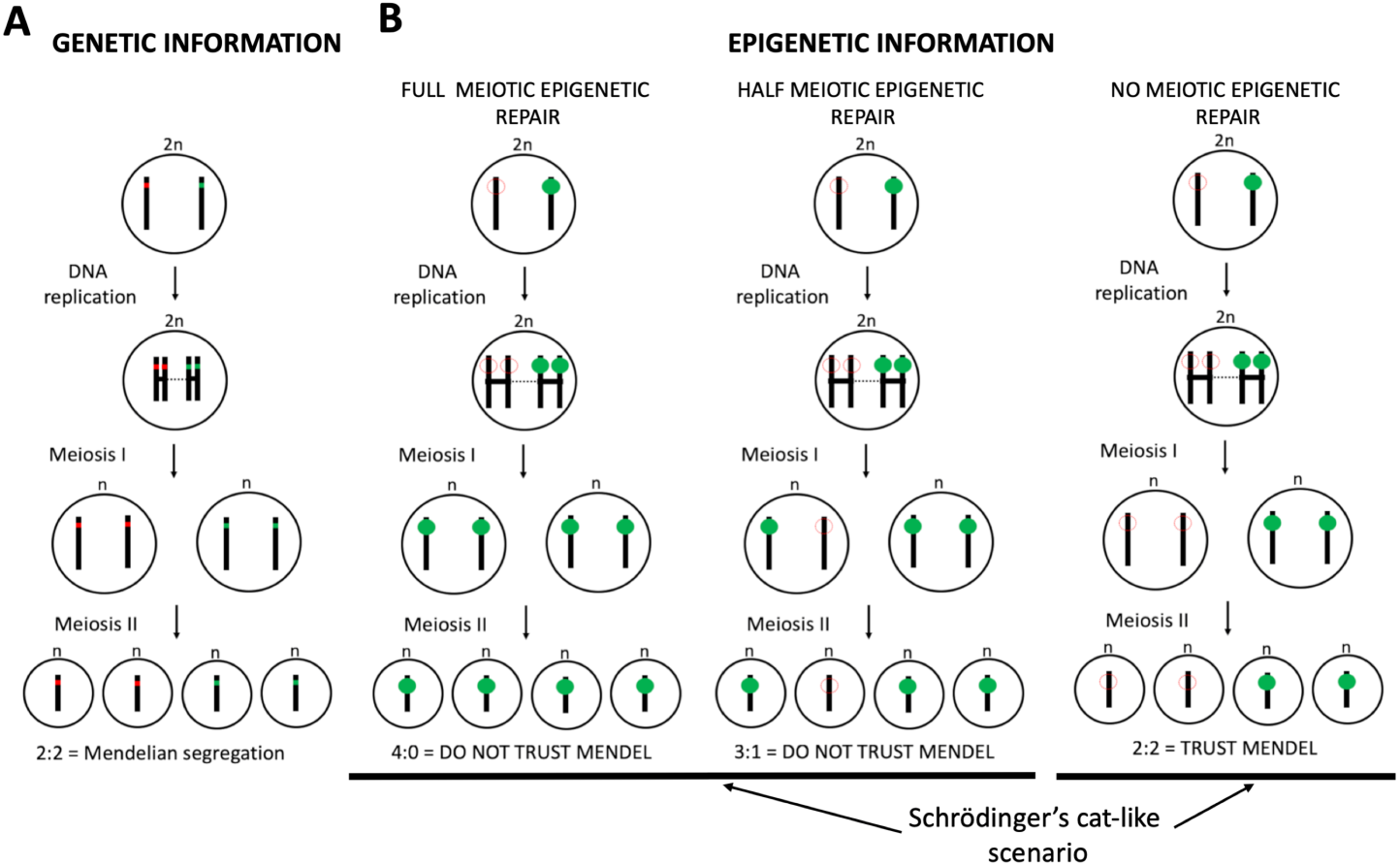
Schrödinger’s cat-like Laws of Mendel for the segregation of the epigenetic information. Schematic representation of a several meiotic processes. **(A)** the Laws of Mendel governing the segregation of the genetic alleles. **(B)** Three panels to outline the segregation of epigenetic alleles. According to the epigenetic repair program hinted in a previous study (Marsellach, 2017), in the examples showing the segregation of the epialleles, three scenarios are contemplated: 1) Full meiotic repair (left), 2) half meiotic repair (middle), and 3) no meiotic repair (right), of a given epiallele (see main text for details). Cells are represented as big circles, one pair of independent solid black lines inside those circles represents one pair of homologous chromosomes (just one pair for simplification purposes). Sister chromatids, once DNA has been replicated, are plotted as joined solid black lines. Pairs of homologous chromosomes are joined by a dotted black line. Genetic DNA alleles are represented as red (defective) or green (wild type) dots, inside the solid black lines (to show that they are part of the DNA fibre). Epigenetic factors (see main text for definition) are plotted as small green circles. The absence of a given epigenetic factor in one locus are plotted as red dotted circle lines instead of the epigenetic factor. Both epialleles are plotted on top of the solid black line (the chromosome), to show that they do not strictly form part of the DNA fibre.

In a previous manuscript I proposed the existence of a pervasive meiotic epigenetic repair program happening in all meiotic cells. In that manuscript, it was as well hinted that: 1) the repair capacity of the cells seems to be limited (probably due to cell resources limitations), and therefore this epigenetic meiotic repair program does not always acts in all *loci*, and 2) there seems to be correlation between this repair capabilities and cell fitness or event lethality avoidance (so that the epigenetic defects are the main cause of cell lethality; see (Marsellach, 2017)).

For the purpose of this standalone paper I would like to ask you to give these hints as given facts. So, I ask you to take the following premises as facts: A) the epigenetic genome-wide reorganization that happens during meiosis (Hackett et al., 2012) brings back previously lost cell functionalities (repairs accumulated epimutations); B) in most cases, in just one meiotic cycle, not all the epimutations are repaired; and C) the epigenetic defects, repaired during meiosis, would ultimately produce cell lethality if not repaired. This implies that the main cause of lethality in mitotic cells are those epigenetic defects.

The above proposed premises imply that the repair of a given *locus* happens randomly (in a limiting scenario some are repaired and some not, which ones is what happens randomly). This makes that some given epimutations are repaired, while some other are not. In accordance with this, it is predicted that, in cells with lots of epimutations a higher number of them would not be repaired; while, in the other way around, in cells with few epimutations it is more likely that all of them are repaired. This describes the ideal scenario named before: a cell with few epimutations entering meiosis would have most of them repaired. Therefore, for the phenotypic characteristic, coded by those particular epigenetic factors (see below for definition of an epigenetic factor), a non-Mendelian ratio will be observed (the Laws of Mendel are not true; see Figure 1B left and middle panels). However, if a cell with lots of epimutations do enter meiosis, then, in some given epimutations repair would not happen. For the phenotypic characteristics coded by these given *epiloci* a Mendelian ratio will be observed (the Laws of Mendel are true; see Figure 1B right panel). In Figure 1B, two scenarios are contemplated for a meiotic epigenetic repair: full meiotic epigenetic repair (Figure 1B left panel), and half meiotic epigenetic repair (Figure 1B middle panel). This is done to contemplate all theoretically possible scenarios. No experimental evidence has yet been obtained so far by this author, that could show that this is the case, further experimental work is needed to test this possibility.

In summary, the randomness nature of the epigenetic repair program happening in all meiotic cells leads to a Schrödinger’s cat-like scenario when studying the segregation of the phenotypic characteristics coded by epigenetic factors.

For the purpose of clarity, I would like to precis what it is referred as an epigenetic factor in this paper. An epigenetic factor is considered any entity that lies “on top” of the DNA fibre, and that, by staying there, affects, somehow, the function of the cell. It is necessary to be pointed that, the alteration of the function of the cell, is not restricted to the alteration of the genetically-coded function of the DNA fibre where the factor is laying, in the case where the DNA fibre bound is a coding region. This definition does include, as well, any modification of the DNA fibre itself that transform, any given nucleotide, in any other nucleotide different than the classical A, C, T or G. Therefore, epigenetic factors include, but does not restrict to: histone modifications, DNA methylation and similar modifications (in organisms where these modifications are present), or any protein, or group of proteins, that binds to the DNA fibre or stays close to it (i.e. a protein not binding the DNA fibre itself, but that binds to another protein binding the DNA fibre).

What I would like to emphasize with this clarification is that, although lots of epigenetic factors have been described so far (Allis and Jenuwein, 2016; Chen et al., 2017), it could well be that there are some not yet experimentally characterized, but functionally relevant, epigenetic modifications.

## 3. The flow of information

How does the information flow inside living organisms? How about from one generation to the following one?

### Biological Information

In order to answer those questions, one has first to clearly clarify what it is understood as information. Nowadays, when one thinks about information in biology, there is a clear predominance to think in the genetic information (the DNA sequence). In recent years, however, the use of the concept of epigenetic information is increasing, even leading to some studies trying to characterize how this information could be passed from one generation to the following one. This brought us the concept of transgenerational epigenetic inheritance (Prokopuk et al., 2015; Wang et al., 2017; Norouzitallab et al., 2019). In the previous section of this article, I have proposed a model to explain the inheritance of the phenotypic characteristics caused by epigenetic factors, which could give an answer, on how this kind of biological information flows from one generation to the following one.

In my opinion, biological information should not be limited to the DNA sequence. Biological information should be any given biological mechanism that leads to a phenotypic outcome. A phenotypic outcome should be understood as the product of the activity of any biological entity. There are phenotypic outcomes clearly visible: affect macroscopic characteristics of the living organisms (i.e. shape or colour), and some others not so clearly visible (i.e. the modification of histone marks in a histone tail), which need specialized technologies for them to be characterized. Remember that, as discussed above, this might include not yet characterized functions and/or modifications of some biological entities.

To further discuss the concept of biological information, one has to go deeper into the history of both genetics and epigenetics. Some reviews are available for this: among others see the following references: (Goldberg et al., 2007; Portin, 2014; Allis and Jenuwein, 2016; Gayon, 2016).

Since Mendel uncovered the laws of inheritance in hybrids (Mendel, 1866), the question of where and how the biolofical information is stored was an open question in the biological field. The rediscovery of Mendel’s Laws in the early 1900s independently by Hugo de Vries, Carl Correns and Erich von Tschermak, and works made by prominent scientist William Bateson marks the start of genetic studies as the science to study heredity (Bateson et al., 1902).

The first clue about where the biological information could be stored came from the chromosomal theory: in the early 1900s, works by Walter Sutton and Theodor Boveri proposed to consider the chromosomes as the bearers of the Mendelian factors (Sutton, 1903; Boveri, 1904). This theory was finally fused with Mendelian genetics by Thomas Hunt Morgan (Morgan, 1915), leading to the establishment of genetics as a new discipline, separated from other biological disciplines. This had as side effect the development of classical genetics, with an abstract non-chemical concept of the gene, and therefore a decline in the interest in the molecular nature of the genetic material (Deichmann, 2004).

The main consensus in the scientific community about what could be the nature of the genetic material during almost the first half of the XX century was that the genetic material was made of proteins (Deichmann, 2004). Two experiments, though, brought the attention to the DNA as the bearer of the genetic information: First, the paper by Oswald Avery, Colin McLeod and Maclyn McCarty, where purified DNA was shown to be able to transform a non-virulent strain of pneumococcus into a virulent strain (Avery et al., 1944), and second, the paper by Alfred D. Hersey and Martha C. Chase, in which they demonstrated that DNA was responsible for multiplication of bacteriophage (Hershey and CHASE, 1952). However, it was not only after the Watson and Crick proposal of a structure for the DNA, that, the DNA was widely accepted as the source of the genetic information (WATSON and CRICK, 1953a; 1953b). Watson and Crick model gave a coherent explanation on how information was stored: a double helix with twice the information would allow the information to be stored, copied, and transmitted from one ancestor to two or more descendants.

Although DNA was clearly accepted as the source of genetic information, it soon became clear that the specific phenotype of a cell was not only due to its DNA content. In parallel to all the development of the genetics field, the epigenetics field started to develop (Allis and Jenuwein, 2016). Historically, the word “epigenetics” was used to describe events that could not be explained by the genetic principles. Waddington defined epigenetics as “the branch of biology which studies the causal interactions between genes and their products, which bring the phenotype into being” (Waddington, 1942; Goldberg et al., 2007). Even before the Watson and Crick model, some phenomena were clearly away from what it was expected just from the well stablished genetic principles. As examples: position-effect variegation (MULLER and Altenburg, 1930), transposable elements (McCLINTOCK, 1951), X-chromosome inactivation (Lyon, 1961) and imprinting (McGrath and Solter, 1984; Surani et al., 1984). Together with this, the concept of genomic equivalence made clear that the DNA sequence of a given cell was not the only factor governing the phenotype that a given cell shows (GURDON et al., 1958; GURDON, 1962).

Proteins were later shown, as well, to be able to transmit biological information from one organism to another. In 1982, the identification of a protein as the “Infectious Particle” in the scrapie disease, showed this (Prusiner, 1982). Strikingly, the problems that both, characterization of DNA as the “Transforming Principle” and prion proteins as “Infectious Particle” had, were similar. In both cases they had to prove the absence of the “opposite” component: proteins or DNA respectively (Prusiner and McCarty, 2006).

Since the beginning of the questioning of where the biological information was physically stored there has been a dispute between nucleic acids and proteins as the real agents of biological information. Therefore, it was considered that either DNA or proteins were the carriers of biological information. This dispute was clearly won by DNA, since the discovery of the structure of nucleic acids (WATSON and CRICK, 1953a; 1953b). But, what if this dispute was only a human invention? Could it be that both, DNA and proteins, together are the carriers of biological information at the same time? (plus some more components, like modified oligonucleotides). In fact, the chromosomal theory identified the whole chromosome as the carrier of the genetic information (Sutton, 1903; Boveri, 1904). Since chromosomes were made mainly of DNA and proteins and, since soon controversial discoveries about the “Transforming Principle” appeared (Deichmann, 2004), an excluding dispute was initiated between both cellular components. In this paper I propose that this was an artificial dispute, and therefore both proteins and DNA (and all other components of the chromosome) contribute to the transmission of the biological information.

### The flow of information

At cellular level, one can see the transmission of biological information in three main scenarios: 1) the flow of information inside one cell, 2) the flow of information between two cells, and 3) the flow of information between a parental cells and cells from its meiotically derived descendants.

#### The flow of information inside one cell

The flow of information inside one cell follows the well know Central Dogma of molecular biology (Crick, 1970) (see Figure 2A). Note, though, that the definition of the Central Dogma does not account for the epigenetic effects that can affect which information is available in a given cell (by allowing/prompting its “reading” or not). One should, then, take into consideration the effect of epigenetic modifications when talking about the Central Dogma of molecular biology (see Figure 2B). One of the main messages of the Central Dogma is that the information flows from DNA to proteins, and not in an inverted order. This means that you cannot translate the amino-acid sequences of a protein in a nucleotides sequences of DNA (Koonin, 2015). This, though, does not mean that proteins cannot affect back the function that a concrete DNA sequence does in the cell (regardless of its coding capacity). Proteins do it, but not through Watson-Crick based code of information, they do it through a completely different code, the epigenetic information code. The amino-acid sequence of the proteins is not translated back to the DNA sequence, but the function that the proteins do, thanks to this amino-acid sequence, does affect back the functionality of the DNA through the epigenetic code. This means that, indirectly, the DNA sequence does create a new layer of information that reverts back into DNA itself, but without affecting its sequence (Figure 2B).

**Figure 2:**
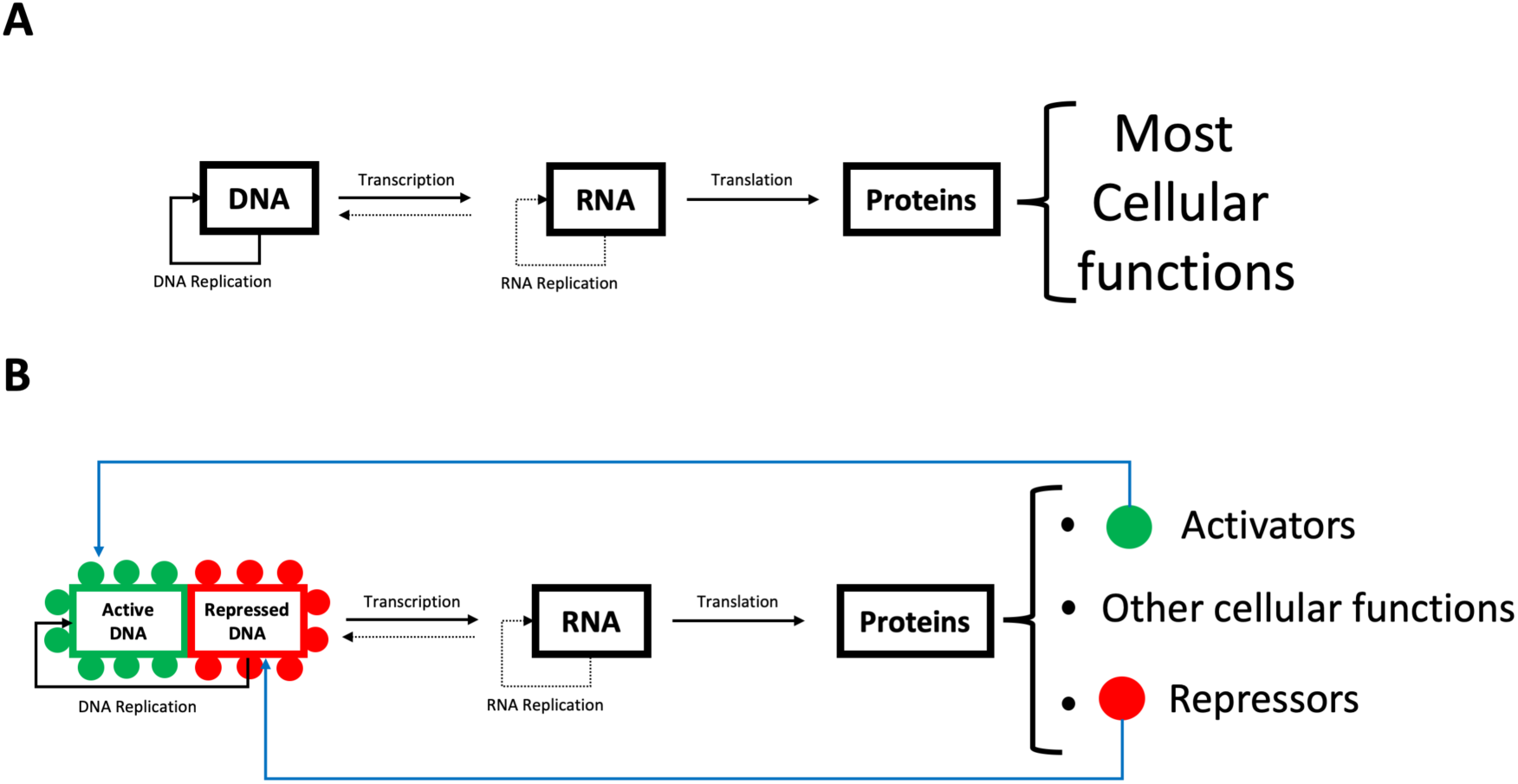
Central Dogma of molecular biology. **(A)** Classical schematisation of the Central Dogma of molecular biology proposed by Francis Crick (Crick, 1970). **(B)** A new schematisation of the Central Dogma of molecular biology which include the effect of epigenetic factors in the flow of information. Note that, as stated in the main text, the feedback carried out by epigenetic factors is shown in blue colour as it does not follow the Watson-Crick based code of information (see main text). Black lines stand for Watson-Crick based information. Solid lines stand for main pathways of information flow. Dotted lines stand for less frequent alternatives. As stated, blue lines stand for epigenetically-coded information, but note that, to plot for the epigenetic code, a single line has been used for the purpose of clarity. To be more accurate many differentiated lines should have been put instead of a single blue line (see main text for details).

I would like to note that the epigenetic code is used by many different players in the cell. Each of those players use a different code (i.e. histone modifications or DNA methylations are two different codes, both of them epigenetic). The only characteristic that they share is that they are “on top” of the DNA fibre, but not part of the DNA fibre itself. In the Figure 2B I have represented with a single blue coloured line the epigenetic code, to differentiate it from the genetic code (represented in black). To be more accurate, though, the blue line should be represented by many lines of different colours (each colour would represent a differentiated code). As discussed earlier, though, we might not even know how many lines we should put on that figure (see Figure 2B). Nevertheless, when referring to epigenetic mechanisms we should name them as epigenetic codes, to highlight that there is not just one code.

#### The flow of information between two cells

The flow of information between two cells has three main different scenarios: A) the flow of information between two “unrelated” cells (see Figure 3A), B) the flow of information between one cell and its identical mitotic descendant (see Figure 3B), and, C) the flow of information between one cell and its mitotically derived differentiated cell (Figure 3C).

**Figure 3:**
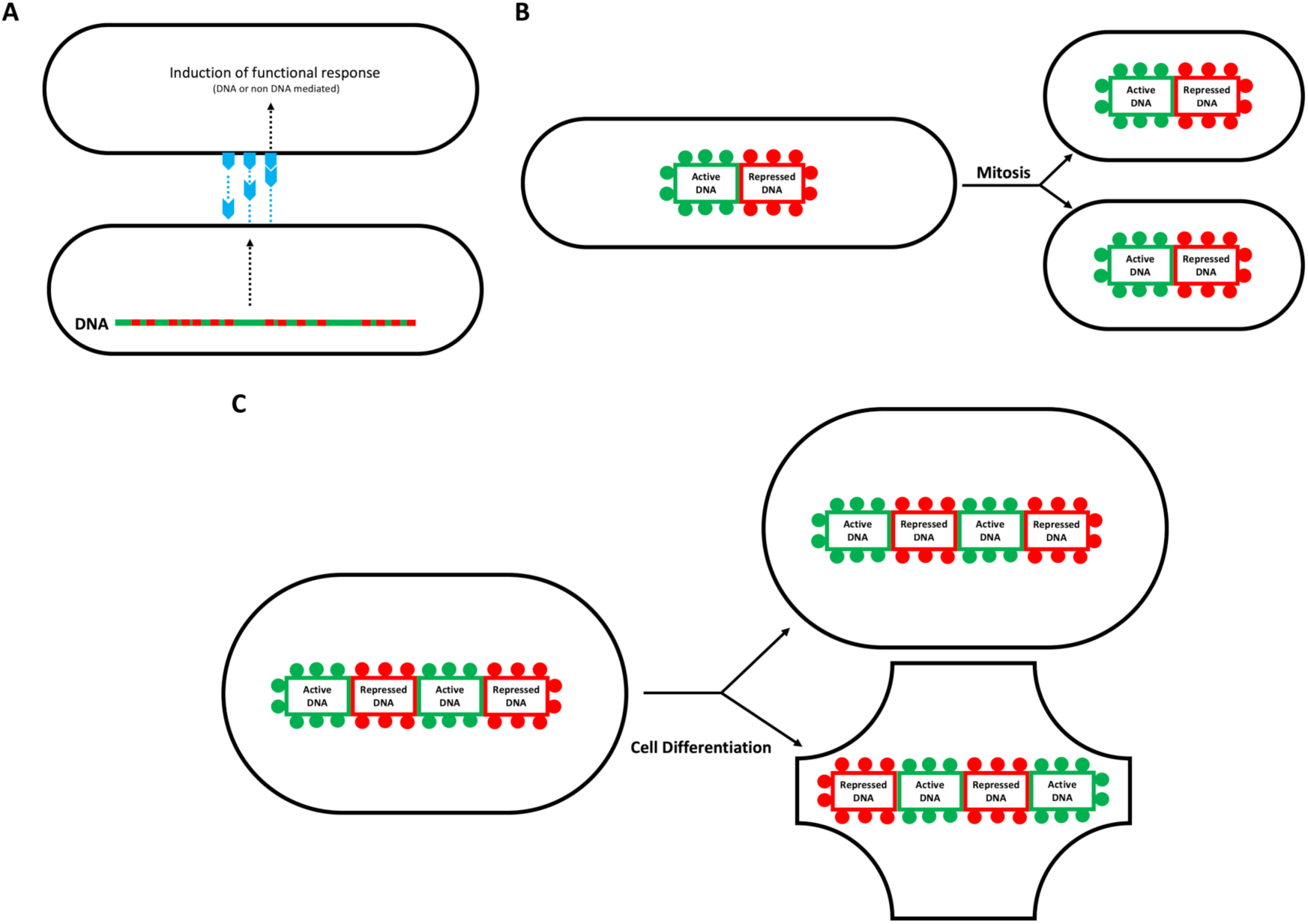
The flow of information between two cells. **(A)** Flow of information between two “unrelated” cells. **(B)** Flow of information between one cell and its identical mitotic descendant. **(C)** Flow of information between one cell and its mitotically derived differentiated cell. In this representation one of the daughter cells keep staying undifferentiated (as the parental cell), while the other differentiate by “reading” different parts of the genetic code (DNA), thanks to the change in epigenetic modifications present in that cell. In **(A)** green line with red dots represents a DNA fibre (green, active regions; red, silenced regions). Black dotted lines represent a bunch of intracellular processes in each of the cells plotted. Extracellular products (secreted by lower cell in the extracellular matrix) and cell surface ligands (three plotted in the cell membrane of the upper cell) and its paths are plotted in soft blue.

It is not the purpose of this paper to discuss about neither cell signalling, mitotic maintenance of epigenetic marks, or cellular differentiation issues. To know more about those topics, the reader is refereed to several reviews and textbooks. Among others in the following references: (Bradshaw and Dennis, 2009; Boland et al., 2014; Almouzni and Cedar, 2016; Atlasi and Stunnenberg, 2017; Palozola et al., 2018)

#### The flow of information between a parental cell and its meiotically derived descendants

The discussion about what might imply the flow of information between a parental cell and cells from its meiotically derived descendants deserves a full section to discuss about it. Therefore, I would like to refer the reader to carefully reads the next section, with the mind open to new ideas.

## 4. The information-based nature of the ageing process

The first and the third premises that I ask you to take as a fact have an astonishing consequence: allow to describe the ageing process as an epigenetic cyclical prosses (see Figure 5). The fact that the main cause of the lethality of the cells are epigenetic defects, and that those defects are corrected during meiosis, allows to explain the ageing phenomenon simply as an access to the information issue. According to these premises, just by controlling which epigenetic information goes through meiosis from one generation to the following one (and, therefore, by explaining the flow of information between a parental cell and cells from its meiotically derived descendants) one can explain the appearance of the ageing process during evolution (see below).

**Figure 4:**
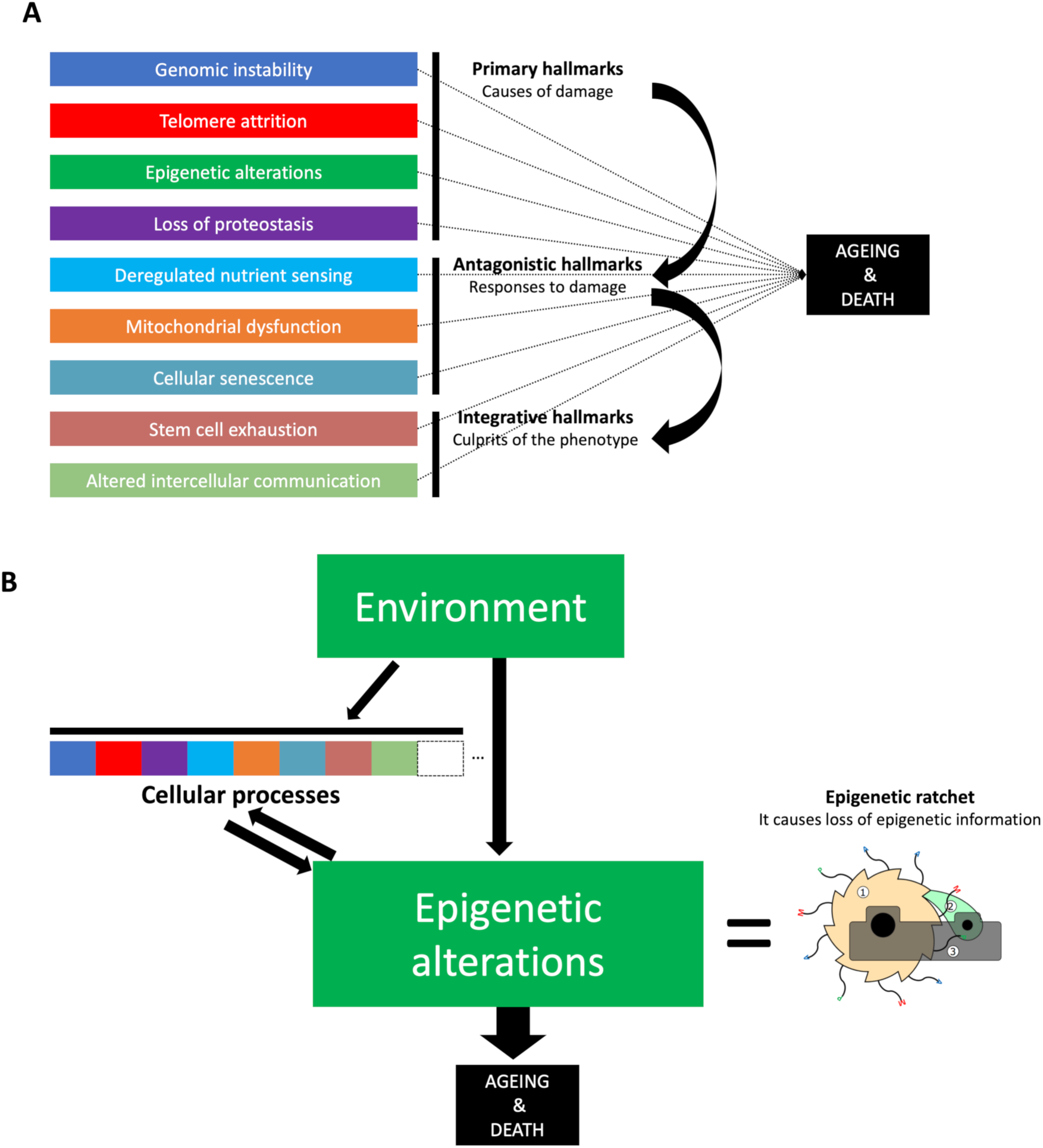
Schematization of the causes of Ageing. **(A)** Adaptation of the schema proposed by (López-Otín et al., 2013). A functional characterization of nine different hallmarks have been proposed. All nine hallmarks have been proposed to influence the ageing phenotype (shown in the figure by dotted black arrows). **(B)** A unique cause of ageing is proposed in this paper. From the nine hallmarks proposed to contribute to ageing (López-Otín et al., 2013), I am proposing that just one, the epigenetic alterations, are the key driver of ageing (See details in the main text).

**Figure 5:**
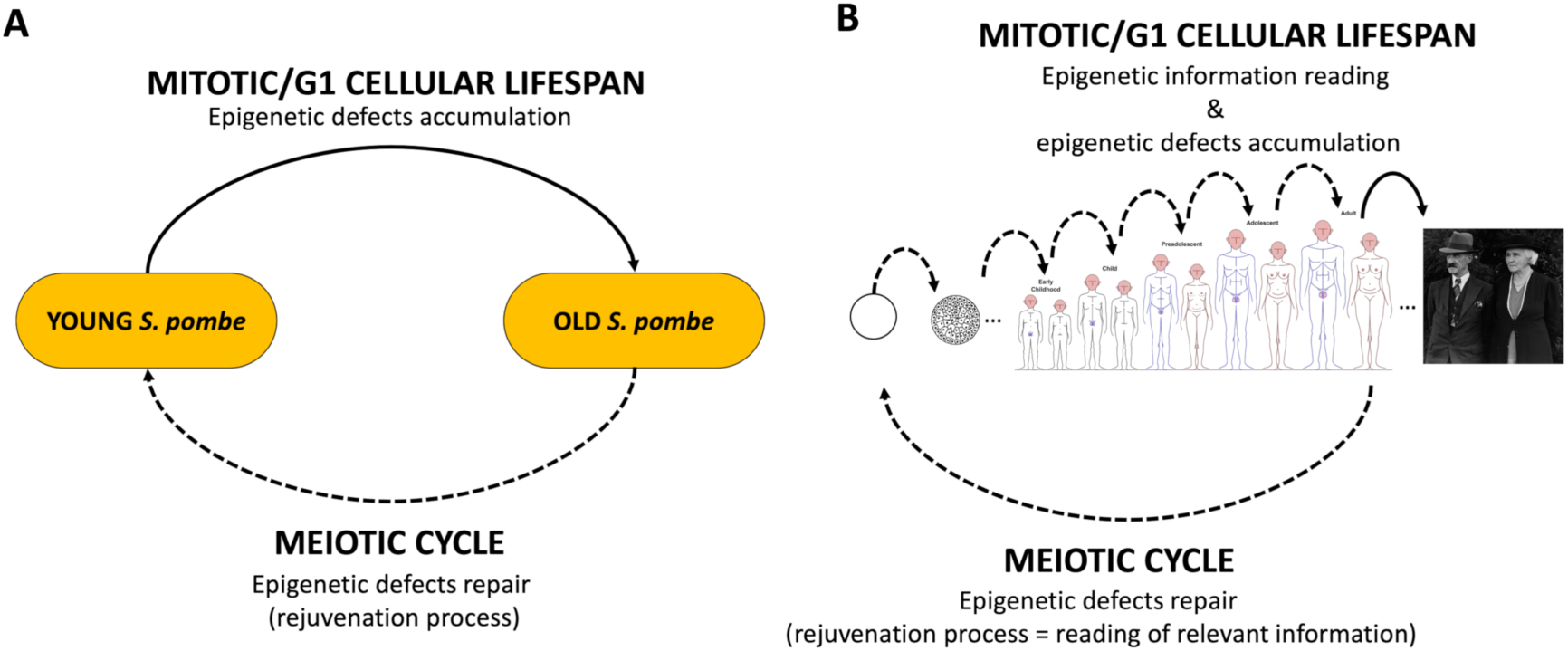
The information-based nature of the ageing process. **(A)** Schematic representation on how a simple cyclical process, with two different phases, can allow explaining the ageing process: 1) the mitotic/G1 phase (top), in which epigenetic defects are accumulated; and 2) a meiotic phase (bottom), in which epigenetic defects are specifically repaired. **(B)** In contrast to what is seen in **(A)**, in multicellular organisms many more steps of “reading of information” are needed to achieve a full functional organism. Dotted lines show steps where “relevant information” is being read to “avoid” ageing. Solid black lines show steps where there is much more losing of epigenetic information that “repair-reading” of it. Drawings used in this figure are adapted from (Marsellach, 2017) or obtained from Wikimedia Commons. Drawings from Wikimedia Commons include: own work by Rajendra Prabhune (human development figures), drawings of Dian∼itwiki (draw of a Morula), and Auckland Museum picture (photo of old gentleman and old lady). All drawings and pictures are distributed under CC BY-SA 4.0 license.

### Ageing is NOT genetically controlled and NOT the consequence of any kind of damage

The discovery of ageing genes: see among others (Kenyon et al., 1993; Kimura et al., 1997; Lin et al., 1997; Ogg et al., 1997; Sinclair and Guarente, 1997; Clancy et al., 2001; Tatar et al., 2001; Kaeberlein et al., 2005), together with the observation that, in distantly related organisms, the effects in longevity of those genes are conserved it generated a large number of studies trying to characterize more and more ageing genes, and the pathways to which those genes contribute. This has led to a flourishing field, in the scientific research community, that aims to characterize more and more of those phenomena, in order to understand why we age. The underlying assumption in this approach is that ageing is the consequence of the action of all those genes and pathways. To date nine hallmarks of ageing have been proposed (López-Otín et al., 2013). Ageing is thought to be the consequence of multiple players, that each individually add its part into the ageing process (see Figure 4A).

On the other hand, although the association of ageing and epigenetics is not new, in recent years there has been an increasing amount of evidences that imply epigenetics as the main actor in ageing (Horvath, 2013; Zhang et al., 2015; Ocampo et al., 2016; Sen et al., 2016; Ashapkin et al., 2017; Maierhofer et al., 2017; Marsellach, 2017), even proposing epigenetics as the only cause of ageing (Gibbs, 2014; Ocampo et al., 2016; Marsellach, 2017). Nevertheless, to date there has not been any coherent explanation on how variability in the epigenome can drive ageing. In this paper I propose that ageing appeared as the consequence of the appearance of the epigenetic codes. To be more precise though, ageing is not only caused by the existence of epigenetic codes, but by how those are dealt by the organisms. In this sense, rejuvenation is a specialized form of cellular differentiation (see text above and Figure 3C) that happens during meiosis. At meiosis the information on how to build a young individual is “read” and passed to meiotically produced cells, which will give rise to the next “young” generation (see Figure 5). The manifestation of ageing is, therefore, nothing else but the loss of this information that occurs in the mitotic descendant cells, that emerge from this meiotically derived (and rejuvenated) cell.

Meiosis is the big success of the eukaryotic cell. In one step both processes rejuvenation (reading of the right information to produce young and fit individuals) and recombination are hold together. As discussed, this generates two ratchets: the well-known genetic Muller’s ratchet, and the newly proposed epigenetic ratchet, which together ensure the constant production of newly-born fit individuals. See Figure 4B and (Marsellach, 2017).

In summary, I’m proposing that ageing is not the outcome of multiple players, all contributing to the same phenotype (an undefined aged phenotype), but the results of a unique process, that has as a side effect the loss of epigenetic information (Figure 4B). The fact that the “young” information is only accessible during meiosis (and silenced in all other cells) has as a consequence a slow and progressive loss of epigenetic information in mitotically dividing cells. Multicellular organisms do age as a whole simply because all their own cells, but the germline, are mitotically dividing cells (See Figure 5B and (Marsellach, 2017)).

I would like to point that by stating that “ageing is not genetically controlled” I do not mean that genetic pathways cannot control lifespan. I just want to emphasize that ageing did not arise as the consequence of the action of those genetic pathways. Ageing is something much more embedded in the fundamental biological processes. Ageing is the consequence of the acquisition of new coding mechanism, the epigenetic codes, which allow the cells to have a much more flexibility to generate new phenotypic outcomes, than the ones achieved only with just the genetic code (the DNA sequence). See Figure 2B for a schematic representation of the epigenetic codes.

It is important, as well to mention, that the conception of ageing that I am proposing, does not lie in the concept of damage. Ageing is not the consequence of damaging any component of the cell, but simply the loss of the epigenetically-coded information that the organisms need to have in order to survive. Damage, though, might appear as ageing progress, but this would be the consequence of the loss of the proper information on how to achieve phenotypic outcomes without damaging cellular components. Damage is the consequence of ageing, and not the cause of it.

The conception of ageing that I am proposing does not need to explain why the germline does not get damaged during ageing (Maklakov and Immler, 2016). Germline derived cells avoid ageing by having access to the information on how to avoid it.

### The “function” of epigenetics

Epigenetics is a word with a complex origin (Deans and Maggert, 2015). Claiming, as I do in this paper, that ageing is epigenetics might prompt to think the other way around: epigenetics is ageing. This statement is in clear conflict with many studies that conclude the involvement of epigenetics in many aspects of cellular and/or organismal functions, other than ageing. I would like to point out that this is just an apparent contradiction. Ageing is not a function *per se*, ageing is, according to my proposal, a loss of information problem. Information that is written using epigenetic codes, and that is used to perform many cellular functions. Losing the knowledge of the many cellular functions written with epigenetic codes, causes ageing to appear. Therefore, I see no problem by claiming in both directions: ageing is epigenetics and epigenetics is ageing.

In summary, the appearance, during evolution, of the epigenetic codes, allowed the cells to perform many more functional outcomes that were not available before. The fact that some epigenetic information is passed to the next generation as a maternal or paternal effect (see later in Conclusions) creates the ageing phenomena, which is crucial for the appearance of complex forms of life (see below). In unicellular organisms, like *S. pombe*, the relationship between epigenetics and ageing, is easier to detect (hints have been observed by this author (Marsellach, 2017)), while in multicellular organisms, which need of the development process in order to be produced, this gets masked by the large number of cellular and/or organismal functions carried out via epigenetic mechanisms (see Figure 5B).

### The membrane-based Object-Oriented-Programming eukaryotic cell

As stated before, meiosis is one of the biggest success of the eukaryotic cell (see above). However, this might not be all: by definition eukaryotic cells are characterized by the presence of a nucleus surrounded by a nuclear membrane. The nuclear membrane might as well be a crucial achievement for its success.

One thing that puzzles ageing research is how easily an organism gets a progeroid syndrome just by the appearance of a single point mutation in the LMNA gene, which codes for lamin A and lamin C proteins (Eriksson et al., 2003; Mounkes et al., 2003). This observation is really difficult to explain by a multiple-players-based origin of ageing. How could it be that the damage that the cells normally accumulate during the whole lifespan of an organism, do appear in very little time, and just by a single point mutation? There is no clear answer to that question with such a conception of ageing. However, conceiving ageing as a loss of information phenomena, does easily allow to propose a mechanism to explain it: the nuclear membrane, together with the proteins that form part of it, are an essential component to maintain the epigenetic information (or the usability of this information). According to this hypothesis, accelerated ageing would appear due to a single point mutation of the LMNA gene by the fact that this would cause a sudden loss of information. In a non-mutant scenario the same information lost would take a lot longer to happen.

As pointed out before (Marsellach, 2017), the acquisition of the epigenetic codes by the eukaryotic cell (on top of the genetic code) is comparable with the transition that happened in human computing by going from just procedural programming languages (the genetic information) to object-oriented programming languages (the epigenetic information). The nuclear membrane might serve as backbone where the epigenetic information is anchored and preserved. Chromatin and 3D nuclear architecture has been shown to be altered during ageing (Zhang et al., 2015; Sun et al., 2018). At the same time, there is a high correlation of epigenetic marks and ageing in many organism (Horvath, 2013; De Paoli-Iseppi et al., 2017). The progressive loss of epigenetic information, that happen during ageing, could lead to an increased nuclear 3D alteration, and to the consequent alteration of many cellular functions. In physiological conditions, reprograming during meiosis might recover the information loss during ageing (see Figure 5 and (Marsellach, 2017)), and restart the “ageing clock”. It has been shown that partial reprograming thanks to Yamanaka factors could improve nuclear envelope architecture and ameliorate ageing related phenotypes (Ocampo et al., 2016). All these observations allow to propose the nuclear membrane as a key evolutionary achievement, that allows the eukaryotic cell to efficiently use the epigenetic codes to regulate phenotypic plasticity, together with the previously evolved genetic code (the DNA sequence), that is shared with less complex forms of life, as bacteria and archaea.

### The “purpose” of ageing

If ageing is simply an information-based issue, why does exist at all? Why not to have all the information always available and, therefore, not to age? In short, why does ageing exist?

The answer to the question of why ageing exists is quite simple: it works. To be more accurate, the question of why ageing exist can be answered thanks to the fact that ageing appeared randomly during evolution, and that it worked to produce complex individuals that do realize that they age, and that ask themselves why.

The more complex an organism gets, the more achievements it can do, but at the same time, there is an increased danger to die by accident. Let us imagine that a complex amortal organism has been produced randomly by evolution. An amortal organism stands for an organism immune to ageing, an organism that, according to the hypothesis defended in this paper, has always access to all the fully functional information. In such a scenario, the chances that all of those organisms died by accident are quite high, and therefore, the chances that none of them progressed much into the evolutionary tree, are as well high. However, mortal organisms (organisms affected by ageing) do constantly produce newly-born and fit organisms. This can clearly overcome the danger created by the accidentality linked to complexity. Each new organism restarts its chances of dying by accident, and therefore, their mortality allowed them to progress through the evolutionary tree. In short, ageing is the mechanism needed in order to allow complex organisms to avoid the death caused by the accidentality linked to a complex lifestyle. Note that the amortal organisms that made it until today are mainly not quite complex organisms (Martínez, 1998; Petralia et al., 2014).

In summary, ageing is just a consequence of the randomness that drives our lives. We should accept randomness as an essential part of our lives.

### The “teleological obstacle”

The last paragraph of the preceding section describes, in my opinion, what is the most important problem of most of today’s ageing research: the teleological approaches that are hidden in most of its reasonings. It has been proposed that today’s biological research is not devoid of teleological thinking (Ribeiro et al., 2015). In my opinion that is something that the whole society has to tackle sooner than later. The biological research community can lead this process by showing how randomness creates one of the biggest mysteries yet to be solved: ageing.

As stated earlier (see above), most of today’s ageing research has its foundation on the characterization of several “ageing genes” (described in more detail in (Marsellach, 2017)). This leads to a development of a whole field whose main purpose was to understand the pathways in which those genes are involved, in order to understand ageing (Partridge, 2010). In the research carried in most of today’s ageing papers, there is implicit the need to fully understand the ageing pathways, the interactions between them and, with external factors (i.e. the environment) affects them, to understand why ageing exists. In my opinion, this is a clear example of the “teleological obstacle” proposed to hinder future progress in scientific knowledge (Ribeiro et al., 2015). Strikingly, this behaviour is seen in both proponents of a programmed view of ageing and of a non-programmed view of ageing (Vijg and Kennedy, 2016). In my opinion, their biggest mistake would be to reduce ageing research mainly to lifespan control research.

In this paper I propose that ageing evolved randomly, but as a consequence of, at least three major transition in biology: 1) the acquisition of a whole new coding mechanisms besides the previously evolved genetic code: the appearance of the epigenetic codes; 2) the appearance of the meiotic step, which, besides the appearance of the recombination step, includes a specialized way to handle the epigenetic information: the rejuvenation step, which involves the reading of the genetically-coded information, and “translating” it into the epigenetic information to be transmitted to the new generation; and 3) the acquisition of a nuclear membrane that allows to anchor the epigenetic information in an spatially ordered way.

## 5. A new paradigm for Genetics, Genomics and Ageing studies

### The external observer

Genetics started as a science in which an external observer, who has direct access to some phenotypical characteristics, tried to use them to deduce the underlying phenomena that can explain how they flow from one generation to the following ones. Later on, with the birth of the molecular biology, the interest was not only in the heredity of those phenotypic characters, but on how they are mechanistically achieved.

Genetically based characteristics (DNA based) were easily identified thanks to the work of Mendel and followers (Mendel, 1866; Bateson et al., 1902; Morgan, 1915). The beauty of Mendel’s Laws, and its astonishing correspondence with some experimental observations, made genetics a flourishing field with many successes. After a dispute about the nature of the genetic material, the determination of the structure of DNA seemed to close the circle (WATSON and CRICK, 1953a; 1953b). Together with many more achievements in the molecular biology field, this leads to a coherent mechanistic explanation about how the information is written and read inside living organisms.

An external observer, however, has only access to the phenotypic information, with no *a priori* knowledge of the mechanisms by which this is achieved. A phenotypic characteristic that is only genetically-coded, will nicely follow the Laws of Mendel. This is easy to detect in the case of genetically-coded phenotypic characteristic coded in just one *locus*, but much more complicated for more complex genetic scenarios. To date, however, there is no accepted proposal of which pattern of inheritance one should expect for an epigenetically-coded phenotypic characteristic. In fact, the underlying consensus in the scientific community, is that no substantial epigenetic information is transmitted from one generation to the following ones (see below). As stated earlier, though, some attention is starting to flourish with some studies focusing in transgenerational epigenetic inheritance (see above). In my opinion, most of those studies do show as well a clear example of “teleological obstacle” (see above), by approaching those phenomena from a Lamarckian perspective.

The model that I am presenting in this paper (see above), together with the experimental framework that I have developed (Marsellach, 2017), allow to test experimentally the pattern of inheritance expected from an epigenetically coded characteristic. First, it allows to detect, thanks to the Advanced Self-cross Lines (ASLs) experiments (Marsellach, 2017), phenotypic characteristics that are epigenetically coded (see Figure 6). Indeed, surprisingly, a high proportion of epigenetic defects is observed in fission yeast wild isolates (Marsellach, 2017). Second, it allows to precisely experimentally test for the expected ratios that the model predicts. And third, it explains why there is not a clear pattern of inheritance observed in the epigenetically-coded phenotypic characteristics: the Schrödinger’s cat-like interpretation of the Laws of Mendel explains why one gets a blurred view, in the epigenetically-coded phenotypic characteristics coded by just one *locus*, instead of the nice view, that one gets with the one *locus* genetically-coded phenotypic characteristics (see Figure 1). This ultimately is due to the existence of an “epigenetic repair program” during meiosis and afterwards (see Figure 5 and (Marsellach, 2017)).

**Figure 6:**
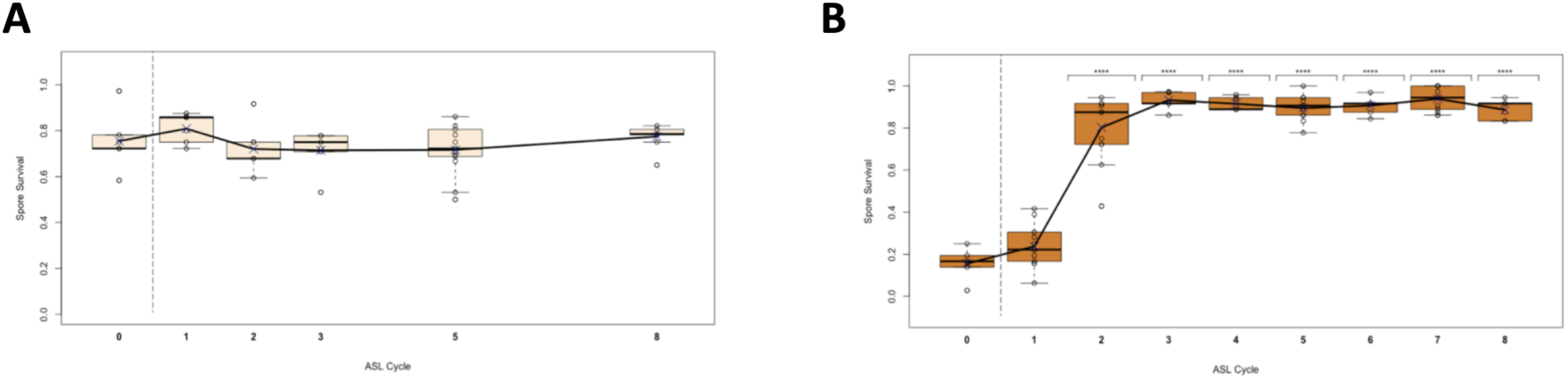
Advanced Self-cross Lines (ASLs). In an ASL cells are self-crossed repeatedly in successive cycles. Parental cells are killed in each cycle, so only new-born spore-derived cells do survive. In a self-cross no genetic variation is expected. At the same time, no substantial retromutations are expected in the average five generations that last each new ASL cycle (Marsellach, 2017). **(A)** Example of an ASL in which no phenotypic change is observed upon eight consecutive meiotic cycles. **(B)** Example of an ASL in which phenotypic change is observed upon eight consecutive meiotic cycles. Figures are adapted from (Marsellach, 2017). For a phenotype genotypically-coded no phenotypic change is expected upon an ASL, as seen in **(A)**, while for a phenotype epigenetically-coded a phenotypic change could be observed, as seen in **(B)** and in many more examples in (Marsellach, 2017).

The model that I am presenting in this paper, though, not only allows that: as a bonus, the same mechanisms that allow to explain for the hidden mechanisms of inheritance of the epigenetically-coded phenotypic characteristics, does offer a coherent explanation for the ageing phenomenon. Ageing is simply a consequence of how the epigenetically-coded phenotypic characteristic are passed from one generation to the following ones (see above and (Marsellach, 2017)).

### Less complex “Complex Traits”?

As stated earlier, the main consensus in the scientific research community is that no substantial epigenetic information is transmitted from one generation to the following ones. This is clearly shown in another flourishing group of papers that approach the inheritance of the phenotypic characteristics as if they were only passed from one generation to the following ones via the genetic code: the GWAS studies (see Figure 7A).

**Figure 7:**
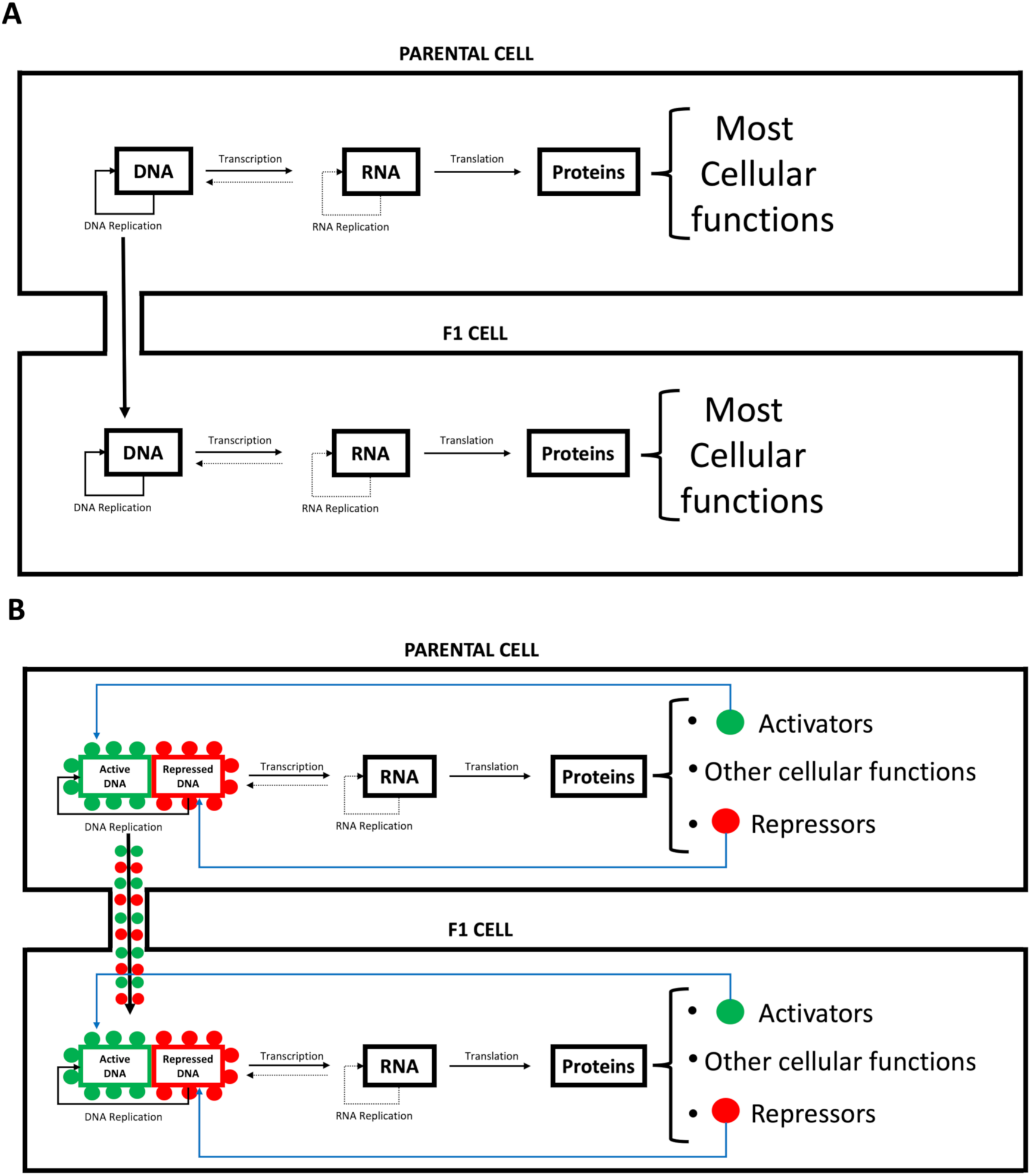
Transgenerational inheritance of epigenetic characters. **(A)** Schematic representation of the main consensus of how information flows from one generation to the following one. **(B)** Schematic representation of the proposed mechanisms of information flow from one generation to the following one. Figure elements are as in Figure 2.

The GWAS studies compare the whole genomic sequences of a group of individuals with the same phenotypic characterization in order to detect which piece of the genetic code is responsible for a given phenotype. This approach, though, has an underlying assumption: all the phenotypic characteristics that one individual shows are derived of its own genetic code. This approach does not take into consideration any other main source of transmissible information from parental cells to its descendants. As explained before this is probably due to the almost unique role that is given to the DNA as the bearer of the inheritable information (see above).

GWAS studies, though, give rise to a central problem nowadays in biology: the missing heritability problem (Manolio et al., 2009). The missing heritability problem is still an open question in the biological field. Many different solutions have been proposed: from the effect of the human microbiome (Sandoval-Motta et al., 2017), to the effect of epigenetic factors (Rakyan et al., 2011). None of those studies, though, question that the DNA is the only bearer of the “genetic” information. The word genetic is put in quotations marks in the preceding sentence because, in fact, the meaning of the word genetic might be a key player of this problem: the word genetic in GWAS is used as exclusively associated with the DNA differences, while, in the classical family studies, the word genetics was referred to the inheritable part of the character transmitted from one generation to the following ones (Bourrat et al., 2017). As explained above, the external observer has no *a priori* information on the mechanistic bases that explains the heredity that he or she are observing. If, as proposed in this paper, the dispute between who is the real bearer of the genetic information was an artificial dispute, and that all the components of the chromosome: the DNA sequence, and all the other components of it (including the many different modalities of epigenetic codes) do contribute to the heritability, this might give a clear answer to the missing heritability problem (see Figure 7B). The missing heritability is clearly not in the DNA fibre as proposed (Bourrat et al., 2017). The missing heritability is linked to the DNA fibre. The missing heritability is caused by all the other components of the chromosome but the genetic code.

All the aspects discussed above have as a consequence that knowing only the DNA sequence of a given individual is not enough to explain all its phenotypic characteristics. One should, as well, know all the information that this individual has coded through epigenetic codes. This might lead to call complex traits to some traits that are no so complex: you call them complex because you cannot explain them, and you cannot explain them, because you limit yourself to just part of the information that would explain them.

In summary, considering just the genetic code, as the GWAS do, and not considering the epigenetic codes, might lead to the impossibility to determine the causative factors that contribute to a given trait.

## 6. Conclusions

The model that I am presenting it simply implies that epigenetically inherited characters have a similar inheritance to the well-known maternal effects characterized in *Drosophila* (Dobzhansky, 1935). However, in this case, it is not as simple as in the previously mentioned example. The main differences are: 1) in the inheritance of epigenetically-coded phenotypic characteristics, both maternal and paternal effects are expected; 2) the inheritance of those characters does not only rely on the previous epigenome of the parental cells/individuals, it relies as well in how they are modified during parental meiosis and during their own developmental processes (see Figure 5 and (Marsellach, 2017)); and 3) the epigenetic codes inherited does not stay stable in the newborns during all their lifespan. As proposed, the epigenome of a given individual might vary due to an epigenetic ratchet acting in all cells (Marsellach, 2017). Indeed, it has been shown that the epigenome does vary along the life of the individuals (Fraga et al., 2005).

This model, although based on preliminary results, is experimentally testable: the ASLs described in my previous work allow the characterization of epigenetically-coded characteristics in *S. pombe*, and probably in other organisms (Marsellach, 2017). The experiments described in that paper, and many more, would allow as well to characterize the degree of implication of epigenetically-coded characteristics into causing ageing and lethality in *S. pombe* cells and other organisms (see Figure 5, Figure 6 and (Marsellach, 2017)).

Another benefit from testing right the model that I am presenting in this paper is that it does not need that Lamarck rises from his grave (Wang et al., 2017). On the contrary, to ensure that no more “teleological obstacles” are present in biological scientific thinking (Ribeiro et al., 2015), Lamarck should be allowed to rest in peace.

## 7. Competing Interests

This author wants someone to hire him in order to be able to experimentally test the model proposed in this article.

## 8. Acknowledgments

I would like to acknowledge David Berman for grammatical and spelling corrections to the first draft of this manuscript.

